# Blood to Biomarker Quantitation in Under One Hour with Rapid Proteomics using a Hyperthermoacidic Protease

**DOI:** 10.1101/2024.06.01.596979

**Authors:** Steven M. Yannone, Vikas Tuteja, Olena Goleva, Donald Y.M. Leung, Aleksandr Stotland, Angel J. Keoseyan, Nathan G. Hendricks, Jennifer E. Van Eyk, Simion Kreimer

## Abstract

Multi-step multi-hour tryptic proteolysis has limited the utility of bottom-up proteomics for cases that require immediate quantitative information. The recently available hyperthermoacidic (HTA) protease “Krakatoa” digests samples in a single 5 to 30-minute step at pH 3 and >80 °C; conditions that disrupt most cells and tissues, denature proteins, and block disulfide reformation. The combination of quick single-step sample preparation with high throughput dual trapping column single analytical column (DTSC) liquid chromatography-mass spectrometry (LC-MS) achieves “Rapid Proteomics” in which the time from sample collection to actionable data is less than 1 hour. The presented development and systematic evaluation of this methodology found reproducible quantitation of over 160 proteins from just 1 microliter of whole blood. Furthermore, the preference of the HTA-protease for intact proteins over peptides allows for sensitive targeted quantitation of the Angiotensin I and II bioactive peptides in under half an hour. With these methods we analyzed serum and plasma from 53 individuals and quantified Angiotensin and proteins that were not detected with trypsin. This assessment of Rapid Proteomics suggests that concentration of circulating protein and peptide biomarkers could be measured in almost real-time by LC-MS.

**TOC Figure:** Rapid proteomics enables near real-time monitoring of circulating blood biomarkers. One microliter of blood is collected every 8 minutes, digested for 20 minutes, and then analyzed by targeted mass spectrometry for 8 minutes. This results in a 30-minute delay with datapoints every 8 minutes.

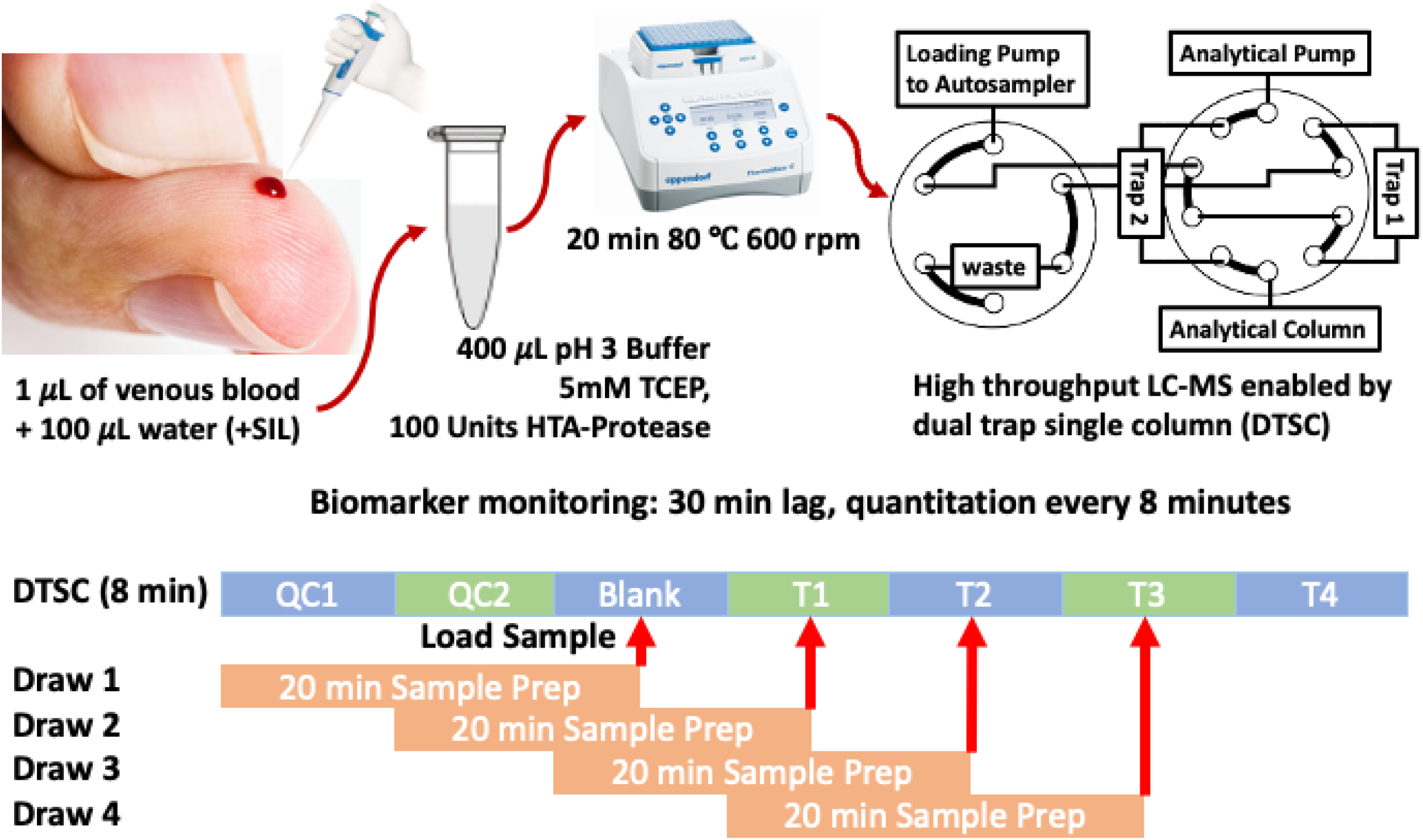

## Introduction

For decades trypsin has been the default protease of bottom-up proteomics due to its efficient and specific cleavage of basic amino acid residues.^1^ Tryptic peptides are long enough for unambiguous assignment to the source proteins, yet short enough to be reproducibly separated with high resolution by liquid chromatography (LC) and then efficiently ionized, transmitted, fragmented, and identified by mass spectrometry (MS). Analysis of LC-MS data of tryptic peptides is aided by the limited search space of predictable tryptic peptides and machine learning algorithms that accurately simulate retention time, ion mobility, and fragmentation spectra from the peptide sequence.^2,3^ Most efficient trypsinization requires reduction and alkylation of cysteine disulfide bridges and denaturation of the protein content prior to proteolysis.^4^ Unfortunately, the reagents used to achieve these prerequisites inhibit trypsin and must be removed or diluted before several hours of trypsinization. Other factors can hamper tryptic efficiency as well.^5^ Innovations in tryptic proteolysis protocols continue to improve on these limitations ^6–8^ but the protocols remain too slow for urgent applications and require training to implement. As such, other proteases are used as alternatives to trypsin alternatives to prepare samples for LC-MS, but are not yet widely adopted.^9^ Here we evaluate the data quality generated by proteolysis of whole blood, plasma, and serum with the recently available hyperthermoacidic (HTA) protease Krakatoa derived from a volcanic hot spring dwelling single-cell Archaea. HTA proteases are remarkably stable and function optimally under conditions that denature most mammalian proteins.

Krakatoa proteolysis is non-specific but preferential cleavage of leucine, aspartic acid, and phenylalanine and large substrates like intact proteins results in reproducible quantitation. Krakatoa is optimally active at a low pH (3.0) and high temperature (>80 °C).^10^ These conditions lyse cells, denature proteins, and provide abundant kinetic energy for rapid cleavage of peptide bonds in just 5-30 minutes. Addition of a reducing reagent to the proteolysis buffer dissociates disulfide bridges, which do not reassemble at low pH so alkylation is not required. Thus, using Krakatoa the entire sample preparation is accomplished in a single swift step. An important benefit of this approach is that short peptides are not cleaved and retain their endogenous form which enables measurement of Angiotensin I and II in blood, serum, and plasma.

Angiotensin I is a 10-residue peptide (DRVYIHPFHL) that is released from Angiotensinogen by renin.^11^Angiotensin I is converted to Angiotensin II by the angiotensin converting enzyme (ACE) through removal of C-terminal leucine and histidine.^12^ Angiotensin II regulates blood pressure by stimulating vasoconstriction. Angiotensin II also stimulates the adrenal gland to release aldosterone which then stimulates the kidney to reabsorb water and salt.^13^ ACE inhibitors impede the conversion of Angiotensin I to Angiotensin II and are recommended as the first line treatment for hypertension.^14^Measurement of Angiotensin I and II and their ratio provides feedback about the efficacy of ACE inhibition so rapid measurement of these peptides may benefit treatment of acute cardiac episodes and other diseases.

Fast proteolysis parallels the current demand in proteomics for high throughput. Modern MS instruments are capable of significantly faster data acquisition than their predecessors,^15,16^ and can generate a deep proteomic profile with short chromatographic gradients of just several minutes.^17,18^ An inefficiency of conventional LC-MS analysis is that no useful data is acquired during sample injection and column cleaning and equilibration. As the gradient is shortened to increase throughput these wasted minutes comprise an increasingly large portion of instrument time. Parallelization of column cleaning and sample loading with analysis of the previous sample results in efficient workflows where data is collected for >80% of instrument operation time.^19–21^

We use the term “Rapid Proteomics” to describe the combination of fast proteolysis with high throughput LC-MS. Rapid Proteomics emphasizes a quick (less than one hour) turn-around from sample collection to actionable data while sacrificing depth of analysis so that MS-based proteomics can be implemented in urgent applications. Potential Rapid Proteomics applications include evaluation of cardiac event severity and risk of subsequent symptoms, quickly observing and mitigating physiological responses to stimuli like medication or allergens, rapidly measuring the severity of toxin exposure, and quick feedback to accelerate iterative optimization of processes. Furthermore, quick and frequent measurement of biological markers approaches real-time monitoring, which could produce a myriad of previously impossible clinical assays. This evaluation found that while Krakatoa proteolysis does not match the depth of trypsin-based proteomics, it is reproducible, quantifies a unique subset of circulating proteins, and enables measurement of bioactive peptides.

## Materials and Methods

All experiments used 1.5 μL packed bed volume Exp 2 bolt cartridge trapping columns (Optimize Technologies) packed with PLRP (Agilent) solid phase and 150 x 0.3 mm analytical columns packed with 2.6 μm Kinetex core shell particles coated with polar capped C18 (Phenomenex). In all experiments chromatographic separations were performed at 15 μL/min flowrate. Electrospray ionization used multi-nozzle M3 emitters inserted into corresponding electrospray sources from Newomics. All Krakatoa proteolysis reactions were carried out at 80 °C in an aqueous buffer containing 5 mM TCEP and 60 mM phosphate citrate at pH 3. All plasma, blood, and serum were obtained under institutional approved IRB (BRANY protocol # HS-2039-528) or purchased (BioReclamation).

### Optimization of Krakatoa Proteolysis Duration

One μL of whole blood collected directly from a finger prick by pipette and 5 μL of pooled human plasma (BioReclamation)^22^ were each diluted to 100 μL with water. Each sample was mixed with 400 μL of proteolysis buffer at 80 °C containing 100 units of Krakatoa (based on manufacturer tyrosine release assay). Aliquots of 100 μL were collected and cooled to 4 °C after 10, 15, 20, 30, and 60 minutes of digestion. Injections of 20 μL from each aliquot were analyzed using a 20-minute Data Independent Acquisition (DIA). LC-MS method on an Exploris 480 MS (ThermoFisher) and Ultimate 3000 LC (ThermoFisher) which was configured for dual-trap single column (**Sup. Fig. 1**).^20,23^

### Reproducibility of Krakatoa Proteolysis

Whole blood from 3 donors (3 μL each) and 5 μL of plasma standard were diluted to 1000 μL with water and separated into 20 μL aliquots that were frozen and stored at -80 °C. Each day, for 3 nonconsecutive days, 5 aliquots of blood from each donor and 5 aliquots of plasma standard were thawed by addition of 100 μL of preheated proteolysis buffer containing 15 units of Krakatoa and digested for 20 minutes. The aliquots were cooled in the autosampler and 30 μL injections were analyzed by 20-minute DIA method on an Exploris 480 using a NeoVanquish LC (ThermoFisher) configured for trap-and elute with trapping column back-flush. The chromatographic gradient, mass spectrometry method and electrospray settings matched the previous experiment.

### Analysis of Atopic Dermatitis and Peanut Allergy Cohorts

Plasma and serum were collected from healthy controls and patients with atopic dermatitis. A history of previous allergic reactions to foods was collected and skin prick testing to a panel of foods (peanut, milk, egg, wheat, soy, shellfish mix (clam, crab, oyster, scallops, and shrimp), almond, English walnut, hazelnut, cashew, Brazil nut, and sesame seed) and common aeroallergens (cat, dog, dust mite, cockroach, mold mix, tree mix, grass mix, weeds mix) was conducted. A subgroup of atopic dermatitis patients was identified as peanut allergic, with a peanut skin prick test wheal≥8 mm, as well as documentation of a previous positive oral food challenge to peanut or convincing history of an immediate allergic reactions to peanut. Both adults and children were included in the study. Written informed consent was provided by the adult study participants. Written informed consent was provided by the parent or legal guardian, and written assent was provided by the child participant, as applicable, before participation. The study was approved by the BRANY Institutional Review Board (protocol # HS-2039-528). The serum and plasma were diluted 5-fold with deionized water, frozen, and stored at -80 °C. To prepare the samples 20 μL of diluted serum and plasma were transferred to 96-well plates, mixed with 100 μL of proteolysis buffer containing 100 units of protease, and digested for 20 minutes.

For comparison, a second set of 20 μL aliquots were digested using an SP3 tryptic protocol^24^ that was automated on a Beckman-Coulter Biomek i7 robot.^25^ Briefly, the samples were dissolved in 2% SDS, reduced for 60 minutes in 40 mM DTT at room temperature, and alkylated with 8 mM iodoacetamide for 30 minutes at room temperature. The proteins were precipitated onto Cytiva Sera-mag beads (ThermoFisher) and washed twice acetonitrile and twice with 80% ethanol. The samples were then digested with trypsin in 50 mM Tris-HCl (pH 8) and 20 mM calcium chloride at 37 °C overnight. The next day, the peptides were eluted off the beads with 0.1% formic acid 2% DMSO.

The Krakatoa and trypsin digested samples were analyzed by a 15-minute DIA-PASEF data acquisition on a timsTOF Pro 2 MS (Bruker) using a dual-trap single column configuration on an Ultimate 3000 LC The Krakatoa samples were also analyzed by DDA-PASEF using an identical separation to generate a spectral library for analysis of the DIA-PASEF (**Sup. Fig. 2**).

### Targeted Quantitation

A set of 10 μL aliquots of 5-fold diluted plasma and serum samples from the peanut allergy cohort were digested by mixing with 90 μL of the proteolysis buffer containing 50 units of Krakatoa and incubating for 20 minutes at 80 °C. The samples were mixed with 100 μL of water containing 10 ng/mL of stable isotope labelled Angiotensin I (1-10 residues), II (1-8 residues), and Angiotensin 1-9. The samples were analyzed using an 8-minute targeted data acquisition (parallel reaction monitoring, PRM) method on an Exploris 480 and Ultimate 3000 using the dual-trap single column configuration (**Sup. Fig. 3**).

### Data Analysis

Spectral libraries for Orbitrap and timsTOF DIA analysis of Krakatoa digested samples were created by compiling data dependent acquisitions (DDA) of the Krakatoa digested samples. The DDA data were analyzed in FragPipe 21.1^26,27^ against the full annotated human database from Uniprot (20,418 proteins) with protease specificity set to nonspecific. The mass error tolerances were set to 10 ppm for precursor and fragment for Orbitrap data and 25 ppm for timsTOF Pro 2 data. The peptide-spectrum matches (PSM) were filtered to 1% false discovery rate (FDR) by target-decoy strategy in the PeptideProphet module, and an FDR cut-off of 30% was applied at the protein level in the first search. The database of proteins identified in the first search was used in the second search instead of the full proteome to reduce the search space. The probable proteins search included deamidation of asparagine and glutamine, oxidation of methionine, peptide C-terminal amidation, and N-terminal pyroglutamine as variable modifications. The reproducibility data was also analyzed with serine, threonine, and tyrosine hexose and phosphorylation as variable post-translational modifications. Peptide spectral matches were filtered to 1% FDR at the peptide and protein levels and compiled into a spectral library using the EasyPQP module. The DIA data were analyzed in DIA-NN 1.8.2^2,28^ using the corresponding libraries with match between runs enabled and quantitation with the QuantUMS module set to “robust LC accurate”. PRM data were analyzed in Skyline-daily.^29^ Each peptide was quantified based on the combined fragment ion peak areas and Angiotensin I and II were quantified as the ratio of endogenous (light) peptide to the spiked SIL peptide (heavy) then multiplied by 50 ng/mL to compensate for the 50-fold dilution during sample preparation and the 1 ng/mL concentration of the spiked standards.

## Results and Discussion

This investigation evaluated the quality of data that is generated by combining fast proteolysis using the Krakatoa protease and high throughput LC-MS. The low pH and high temperature at which the Krakatoa protease is most active are conditions in which most cells lyse and proteins denature. Addition of 5 mM TCEP to the reaction disrupts the disulfide bridges. At pH 3 the disulfide bonds do not re-assemble, as evidenced by the many identified peptides that contained unmodified cysteine. Since no alkylation is required, the entire sample preparation process is completed in one step. The trapping columns used in the LC configuration cleaned and desalted the samples online and parallelized sample injection with analysis of the previous sample. This allowed a seamless transition from a single-step sample preparation to a fast and efficient LC-MS analysis.

### Krakatoa Cleavage-site and Substrate Selectivity

Optimization of proteolysis duration reaffirmed the recently published observation that the Krakatoa protease preferentially cleaves at the C-terminus of leucine, glutamic acid, and phenylalanine (**Fig. 1A**).^10^ Krakatoa reactions also cleaved other residues at a lower frequency and in the limited reaction conditions tested showed missed cleavages, as reflected in the wide distribution of peptide lengths (**Fig. 1B**). Direct comparison of 10, 15, 20, 30, and 60 minutes of Krakatoa proteolysis of whole blood and plasma found that products of 10 minutes of proteolysis increased analytical column back pressure and high charge envelopes were detected during the high organic column flush, indicating undigested protein. The high charge clusters decreased at 15 minutes and were absent with 20 minutes of proteolysis (**Data not shown**). Qualitatively, the same peptides were identified at all time points, but the intensity of the long peptides (>25 residues) decreased after 60 minutes of proteolysis and the intensity of smaller peptides concomitantly increased. The difference in cleavage rate for proteins (not detected after 20 minutes), and long peptides (still present but less abundant after 60 minutes), suggests an additional protease selectivity for larger substrates (**Fig. 1C**).

**Figure 1.**
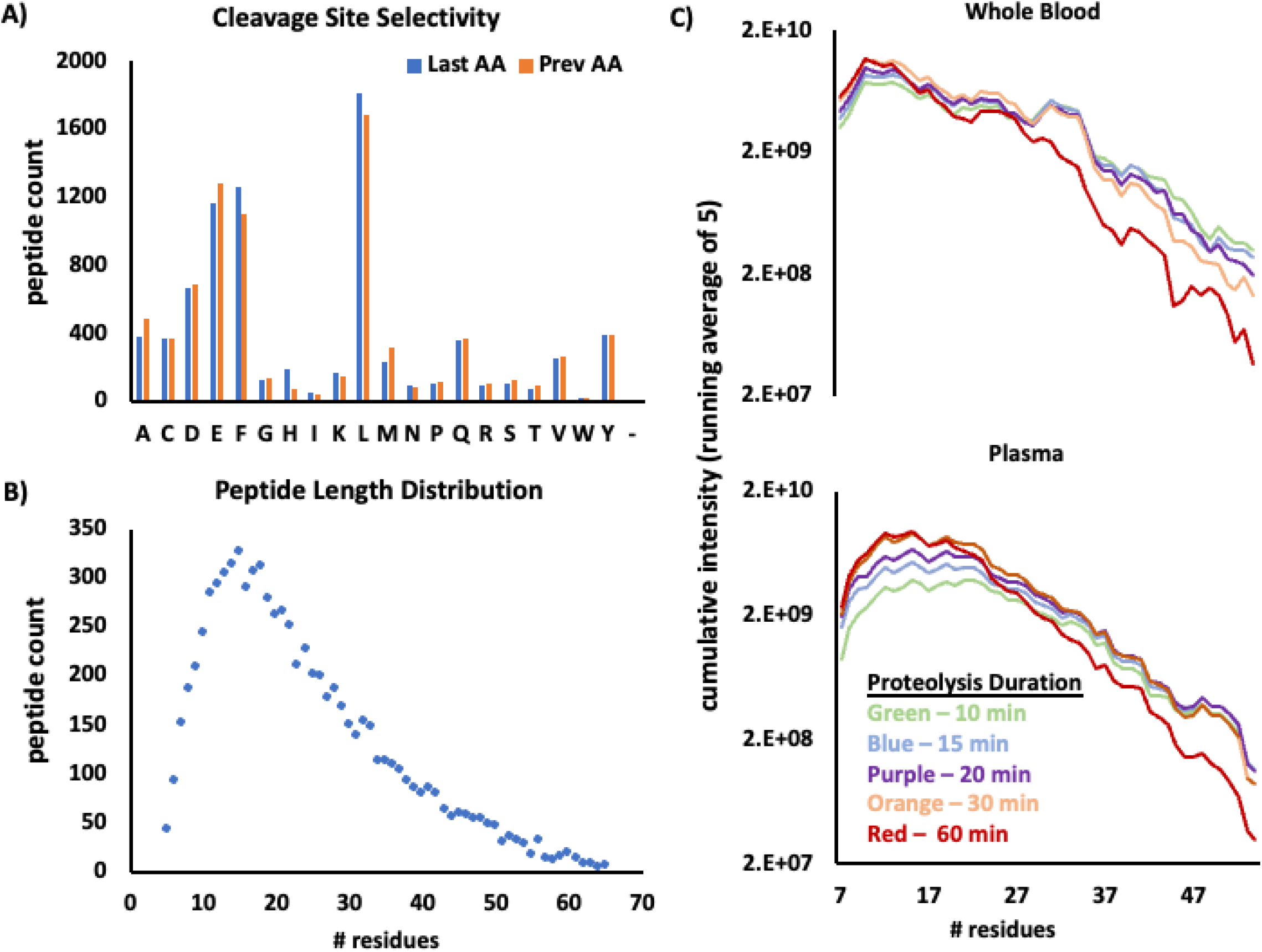
**A** – Distribution of cleavage sites in blood and plasma show that the Krakatoa protease most frequently cleaves at the C-terminal of leucine, phenylalanine, and glutamic acid, but other amino acid residues are cleaved as well. **B** – Distribution of peptide length, while most of the generated peptides are between 10-20 residues, much larger peptides are generated as well, suggesting a high missed cleavage rate. **C** – Average peptide intensity relative to length after different durations of proteolysis, after 60 minutes the biggest peptides diminish while the smaller peptides slightly increase in whole blood (top) and plasma (bottom).

Given these results we set a singular reaction condition for this study: 20 minutes at 80°C in pH 3 60 mM phosphate citrate buffer with 5mM TCEP. These parameters maintain fast turn-around while avoiding undigested protein which would interfere with the chromatographic separation. Sample preparation could potentially be accelerated with increased temperature and/or protease amount. However, 80 °C was used to limit evaporation, and 100 units of Krakatoa protease were used per μL of blood, or 3-5 μL of plasma. These conditions were deemed sufficient to carry out this study and further optimization of HTA-protease dosage and reaction conditions are expected to yield further gains but are beyond the scope of this initial study.

### Evaluation of Reproducibility

To determine the proteolysis quantitative reproducibility, 5 whole blood aliquots from 3 donors, and 5 in-house plasma standard aliquots were digested and analyzed on 3 nonconsecutive days. Over 160 proteins and around 5,000 precursors were quantified at inter-day CV < 30% in all three blood samples. In plasma, 130 proteins and over 5,000 precursors were quantified at inter-day CV <30% (**Fig. 2**). While one might expect that protease specificity is a requirement for reproducible quantitation, other low specificity proteases have also been shown to achieve reproducible quantitation.^30^

**Figure 2.**
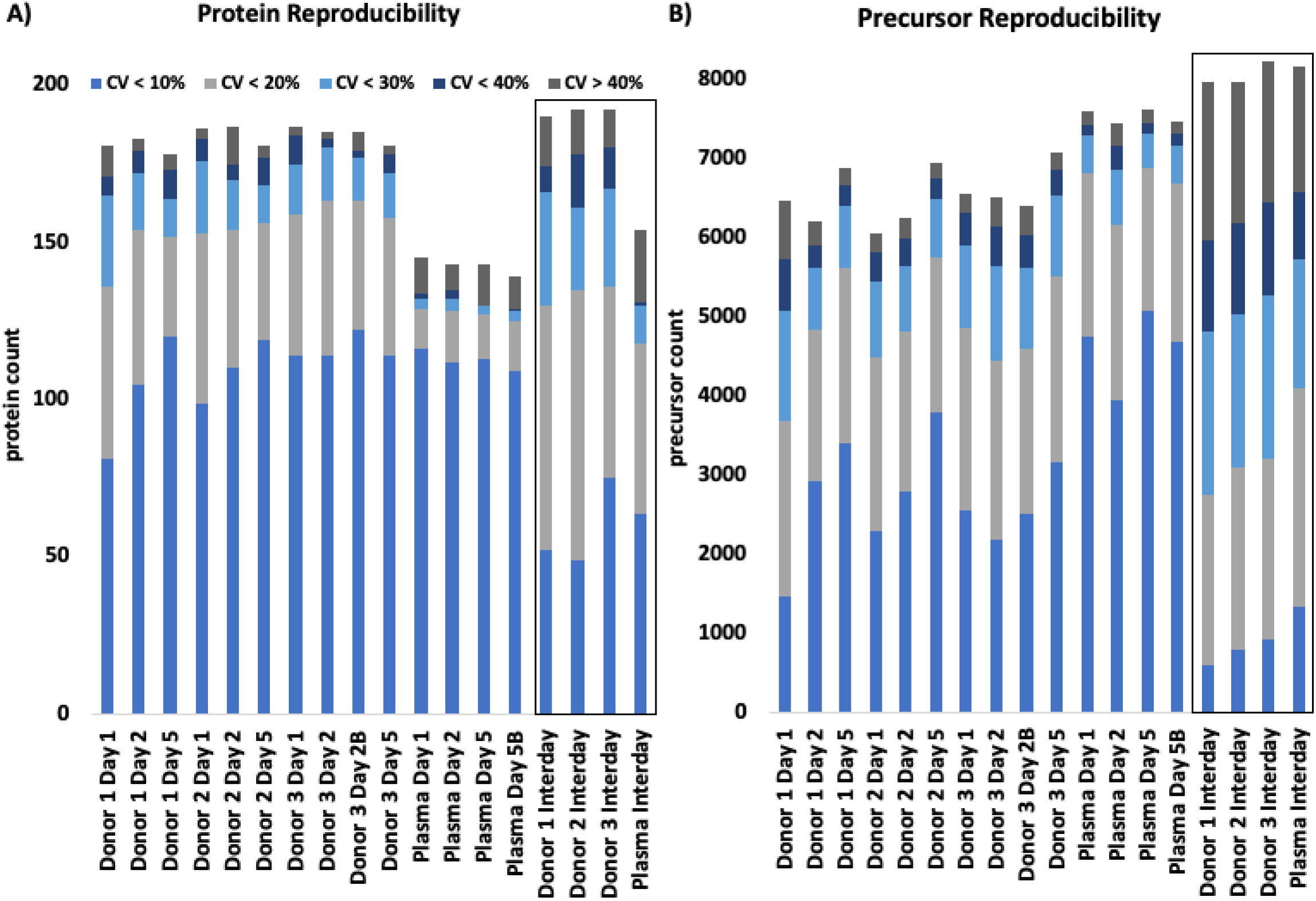
**A** – Reproducibility protein quantitation using five aliquots of whole blood from three donors and a plasma standard on three non-consecutive days along with the inter-day combined reproducibility (outlined with box). Most proteins had a CV <30% (light and medium blue and light grey). **B** – Reproducibility of precursor quantitation as outlined in panel A.

Proteins with inter-day CV <40% and >90% data completeness in the three blood samples were considered reliably quantifiable. Plotting the logarithmically transformed intensities of these 100 proteins across the 15 replicate injections demonstrates that subtle differences in abundances can be observed among healthy controls with this methodology. Analysis of the data which included glycation and phosphorylation found that these post-translational modifications were detectable in Hemoglobin A B, D, and Albumin. The hemoglobin modifications were similar between donors which suggests that there is a stable baseline in healthy blood (**Fig. 3**). This proteomic profile was generated from 20 minutes of sample preparation and 20 minutes of LC-MS analysis, amounting to a less than one hour turnaround.

**Figure 3.**
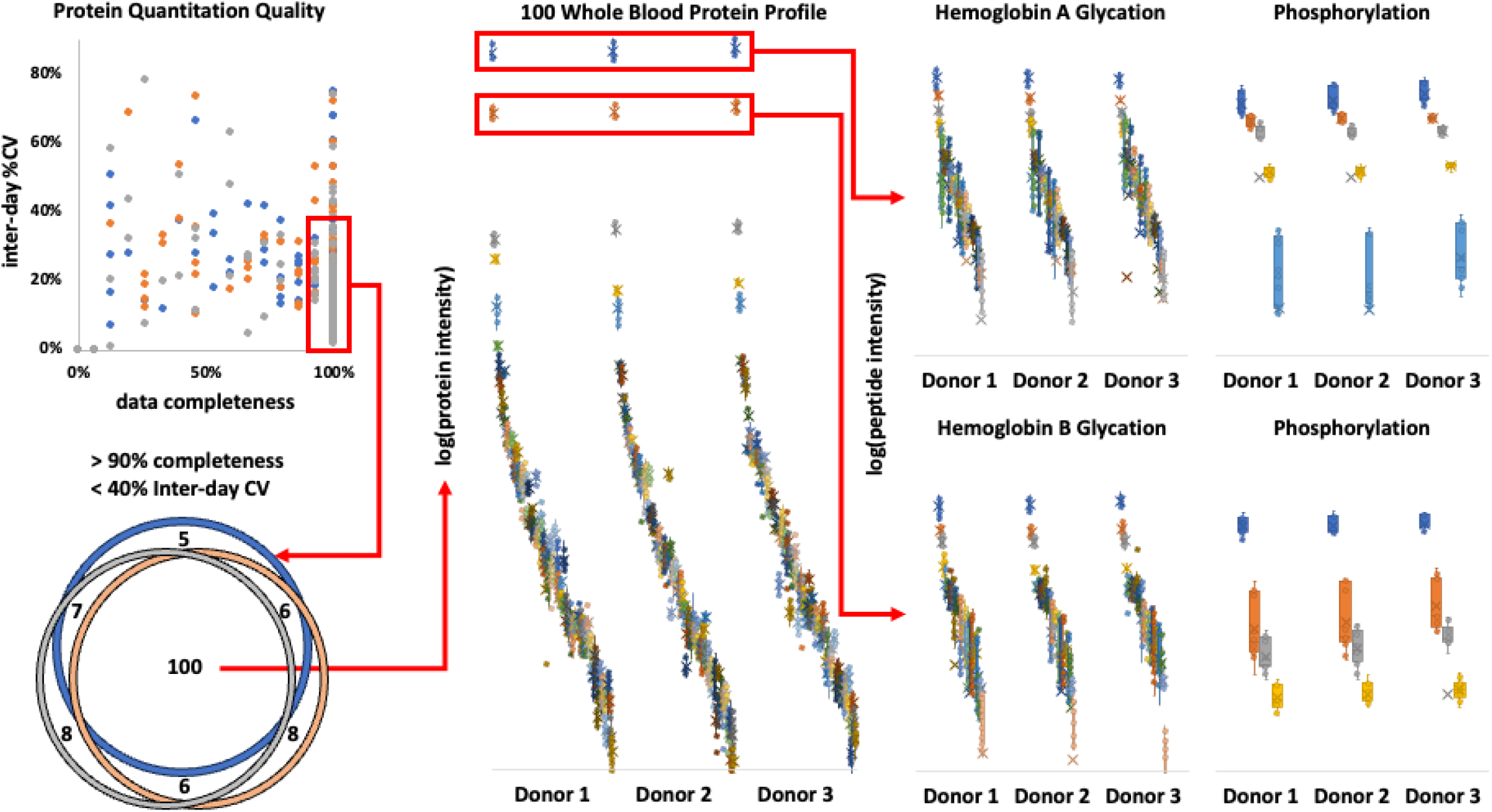
Reliably quantified proteins (CV < 40%, completeness > 90%) in all three donors (scatter plot and Venn diagram) could be used to generate a quick proteomic profile for each donor. The plot of their logarithmically transformed intensities from all 15 reproducibility runs (middle) shows subtle differences for each donor. Peptides from abundant proteins containing post-translational modifications were quantified and appear to be in similar abundance between the three healthy donors (right).

### Analysis of Plasma and Serum of Patients with Atopic Dermatitis and Food Allergy

This methodology was applied to analysis of plasma and serum collected from 18 healthy controls and patients with atopic dermatitis with and without food allergy to peanuts (15 positive and 20 negative). After the 20-minute sample preparation the samples were analyzed by 15-minute DDA and DIA-PASEF on a timsTOF Pro 2. The DDA runs were used to generate a library for DIA analysis. The same cohort was digested with trypsin overnight and analyzed with the same DIA-PASEF method and library-free analysis in DIA-NN 1.8.2 for comparison. The number of precursor identifications was stable and similar for Krakatoa (3,300-3,400 excluding modified peptides) and trypsin (3,300-3,600), however twice as many proteins were identified in the trypsin digested samples (**Fig. 4A**). The reduced sensitivity can be attributed to over-representation of albumin in the Krakatoa digest. In all samples combined over 4,000 albumin peptides (56% of total identifications) of varying lengths redundantly representing the same regions were detected due to nonspecific cleavage. By contrast, trypsin generated only 122 albumin peptides (**Fig. 4B**). Fewer albumin peptides in the tryptic digest reduced ionization suppression and enabled detection and quantitation of peptides from lower abundance proteins.

**Figure 4.**
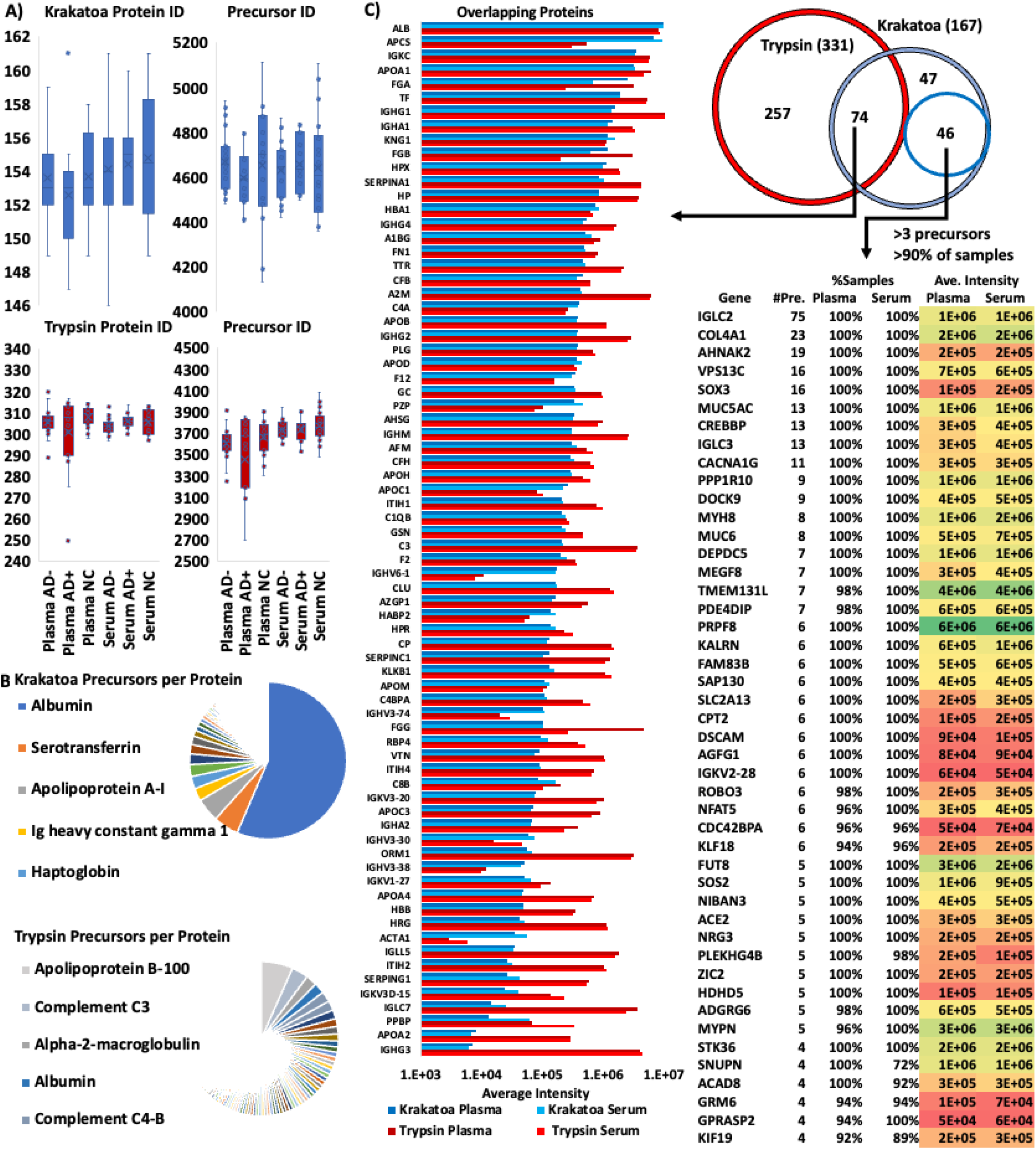
Comparison of analysis of serum and plasma from 53 individuals with Krakatoa and trypsin. **A** – Protein and precursor identifications between trypsin and Krakatoa. Identifications are level between groups, but trypsin identifies twice as many protein species. **B** – The Krakatoa protease identified mostly albumin peptides which reduced sensitivity, while with trypsin the peptides were more evenly distributed. **C** – Comparison of protein identifications between the proteases, only 74 proteins were identified by both and their abundances were differed by protease. 93 proteins were unique to Krakatoa and 46 were detected with >3 precursors in >90% of either plasma or serum samples. NC: normal healthy controls, AD-: patients with atopic dermatitis without food allergy, AD+: patients with atopic dermatitis and peanut allergy.

Control of FDR during generation of the spectral libraries posed a further challenge. Conventional target-decoy based FDR approximation creates a population of decoy peptide sequences equivalent in size to the hypothetical possible peptide population, but with nonspecific proteolysis the number of possible peptides and decoys is many orders of magnitude higher than the real peptides in the sample, which decreases the search sensitivity (number of real peptides identified). We reduced the search space to only probable putative proteins to increase the size of the spectral library from 6,211 to 8,559 peptides. However, the cut-off for “probable” putative proteins was difficult to determine. With no FDR filter at the protein level in the initial search, angiotensinogen was identified but several highly improbable proteins that we considered false positives were incorporated into the spectral library as well. We unambiguously quantified Angiotensin I and II in all of the samples using targeted analysis but Angiotensinogen, the protein from which these peptides are derived, did not meet the 30% FDR threshold that was used for this analysis. Improvement of FDR control and other innovations in data analysis for nonspecific proteolysis would increase depth of analysis.

The overlap between trypsin and Krakatoa sample preparations was only 74 proteins, with the majority of proteins unique to either protease. The differences in average intensities of the overlapping proteins (**Fig. 4C**) indicates that each protease emphasizes different proteins. These quantitative differences between proteases can be attributed to differences in peptide ionization efficiencies, susceptibility to proteolysis, or matrix ion suppression. A total of 93 proteins were quantified by Krakatoa that were not detected with trypsin and 46 unique proteins met the cautious filter of >3 precursors and detection in >90% of either plasma or serum samples (**Fig. 4C**). This filtered list includes some interesting potential biomarkers like CREBBP,^31^ AHNAK2,^32^ SNUPN,^33^ and transcription factor SOX3.^34^ The increased sensitivity for these proteins could be more important in some applications than total number of identifications and Krakatoa offers a complementary perspective to conventional trypsin dependent proteomics.

The quantitative differences between the proteases propagate to protein differentiation between cohorts (**Fig. 5**). Comparison of peanut allergy positive to negative atopic dermatitis patients and normal controls to all atopic dermatitis patients shows that different sets of proteins appear differentiated between the cohorts depending on the protease used. Some proteins identified by both proteases show discrepant differentiation like Transthyretin (TTR) which appears slightly but significantly (*p* < 0.05) elevated in the serum and plasma of the peanut allergy positive cohort in the Krakatoa proteolyzed samples but does not appear differentiated when digested by trypsin. With both proteases the differences between cohorts were subtle, so quantitative differences as well as changes in how the samples are normalized could shift the perspective on protein differentiation. TTR and C1Qb, a protein that appeared to be elevated in plasma and serum of atopic dermatitis patients compared to controls were included in targeted re-analysis of the samples to validate these initial observations.

**Figure 5.**
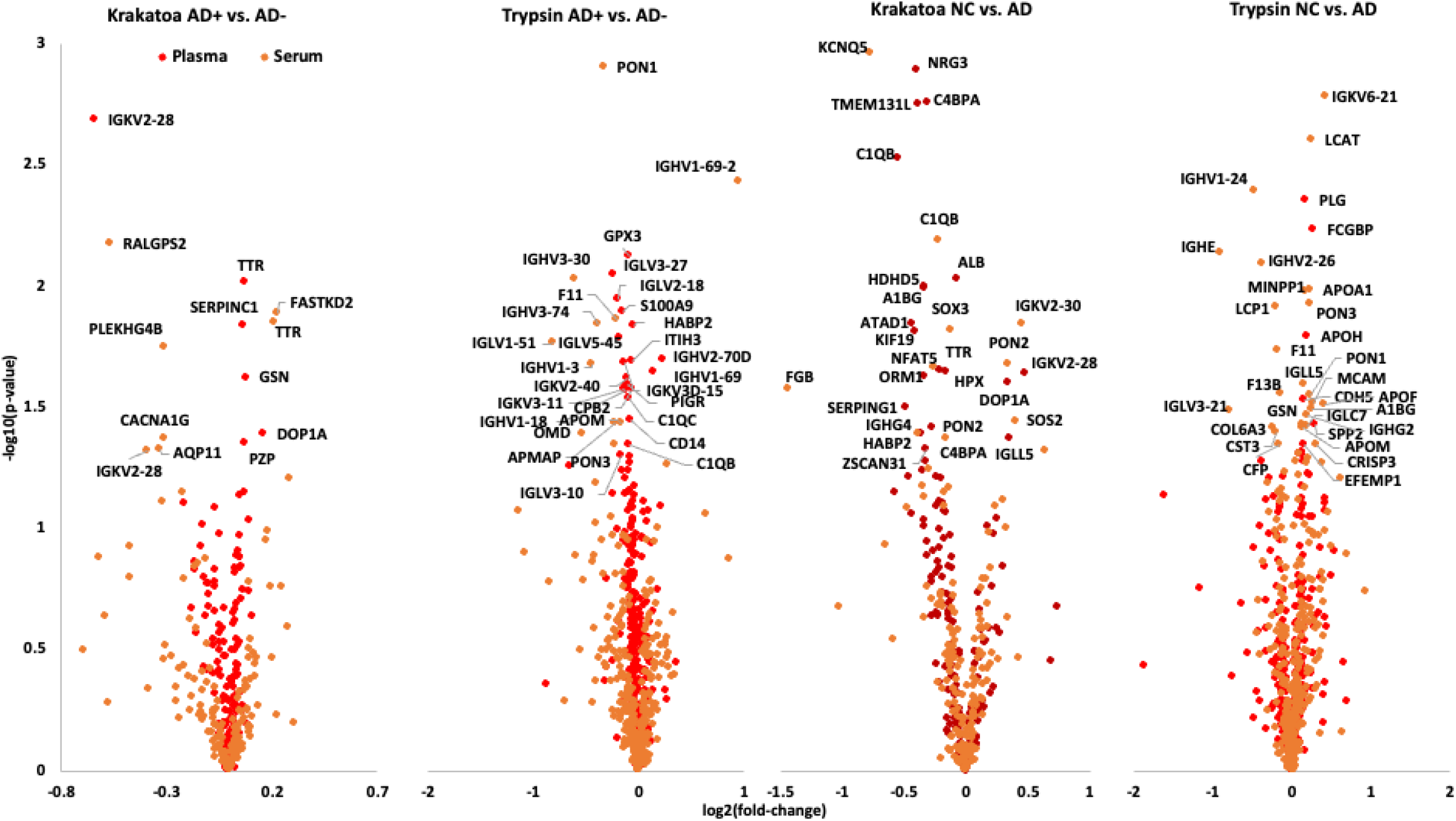
Volcano plots comparing atopic dermatitis patients with (AD+ n = 15) and without (AD-n = 20) peanut allergy and normal controls (NC n = 18) to all atopic dermatitis patients using Krakatoa and trypsin. In all comparisons the changes are subtle (low fold change), but proteins that differentiate (p < 0.05) two groups are labelled.

### Rapid Angiotensin I and II Assay

An 8-minute (injection to injection time) parallel reaction monitoring (PRM) LC-MS method was developed to quantify Angiotensin I, II and 1-9 with matching stable isotope labelled (SIL) standards. The method also included peptides from C1Qb and TTR to demonstrate that this approach can also quantify several proteins along with bioactive peptides, which is not feasible with extraction of bioactive peptides by organic solvent precipitation of the protein content. Targeted quantitation of C1Qb confirmed that the protein was significantly (*p* = 0.0002 for peptide 1 and 0.015 for peptide 2) increased in the plasma of atopic dermatitis patients compared to controls, and significantly (*p* = 0.005, 0.008) decreased in the serum of peanut allergy positive compared to allergy negative atopic dermatitis patients. Only two variants of the same TTR peptide appeared significantly elevated in the plasma of the peanut allergy positive atopic dermatitis group, suggesting that this protein is not really differentiated between cohorts.

The targeted analysis quantified Angiotensin I and II in all samples while also measuring two circulating proteins with a less than 30-minute turn-around. Comparison of serum and plasma Angiotensin for each donor showed a high degree of agreement between the two matrices and range of 3.8 to 79.5 ng/mL for Angiotensin I and 10.6 to 212.2 ng/mL for Angiotensin II (**Fig. 6**). Despite the wide ranges, the ratio between Ang I and Ang II remained at 0.37 +/-0.02 in plasma and 0.38 +/-0.04 serum. The Angiotensin 1-9 variant was included in the assay to check if proteolysis of circulating angiotensinogen could create new Angiotensin peptides. If Angiotensin I and II were generated by proteolysis the middle peptide should also be generated. Angiotensin 1-9 was not detected in any sample and the low signal from other Angiotensinogen peptides suggests that the quantified Angiotensin I and II are endogenous. We did not expect the Angiotensin levels to distinguish between patients with and without peanut allergy and healthy controls, and indeed the variance between individuals far exceeds any differences between groups. However, this dataset demonstrates that the methodology can measure the bioactive peptides across a wide range of concentrations.

**Figure 6.**
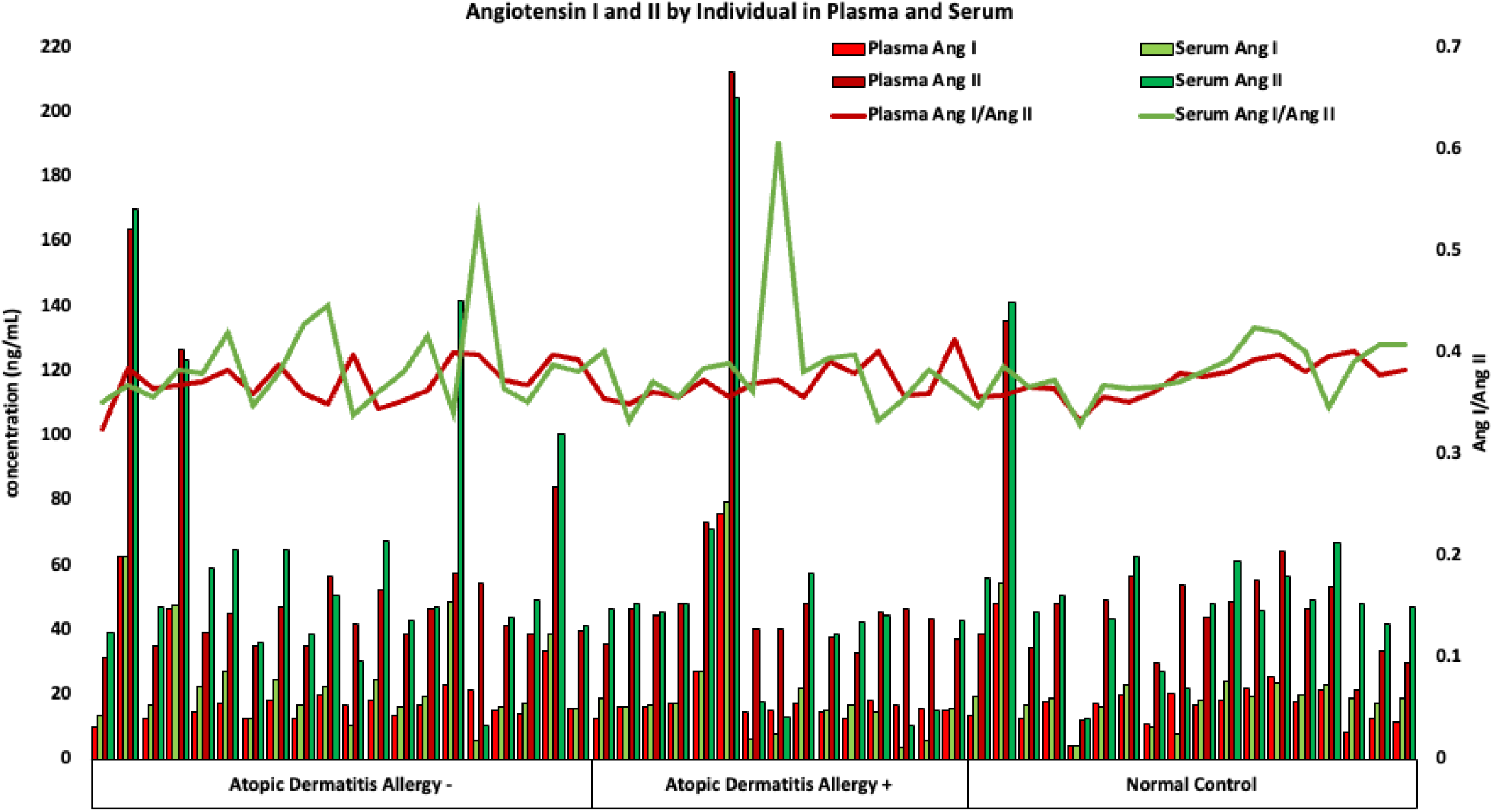
Angiotensin I and II measurements for each donor in serum and plasma. Despite the wide range of concentrations within each cohort the serum and plasma measurements are consistent and the ratio between Ang I and Ang II remains stable. Allergy-: patients without peanut allergy; allergy+: patients with peanut allergy.

### Conclusion

The goal of Rapid Proteomics is to quickly quantify protein and peptide biological markers to inform urgent decisions. The Krakatoa protease digests blood, plasma, and serum in a single 20-minute step. Subsequent high-throughput untargeted LC-MS analysis generates a profile of 100 reliably quantified proteins from 1 microliter of blood in under an hour (20-minute preparation + 20-minute LC-MS analysis). While this approach does not match the depth of trypsin-based sample preparation, it enables measurement of proteins not normally detected by trypsin. The fast proteolysis was also paired with targeted mass spectrometry for a turn-around of under 30 minutes (20-minute preparation + 8-minute LC-MS). This potential was illustrated with targeted quantitation of Angiotensin I and II, two crucial elements of the RAAS, along with two proteins. The panel of complementary proteins can be adapted for specific condition and increased further with a faster mass spectrometer. Other bioactive peptides may be detectable as well. The ability to measure biomarkers with a 30-minute turn around offers an unprecedented opportunity to monitor physiology at a chemical level almost in real-time (**TOC Figure**).

## Acknowledgements

Research reported in this publication was supported by the National Institute of General Medical Sciences of the National Institutes of Health under award number 5R44GM128540 (SMY) National Institute of Allergy and Infectious Diseases grant 1U01AI152037–01 (DYML, EG), Funding from grants R01HL111362 and U01 DK124019-01 (JVE) and Cedars-Sinai Winnick Family Research Center Award (AS) and the Barbara Streisand Women’s Heart Center in the Smidt Heart Institute. JVE is the Erika Glazer Endowed Chair in Women’s Heart Health. We would like to thank Cedars-Sinai Proteomics and Metabolomics Core and Precision Biomarker Laboratories for access to staff and instrumentation. We thank Allison Brand-Narlock and Parker Brand for sharing their insights and for HTA-Protease production and QC, National Jewish Hospital for providing human plasma and serum samples, and Laurie Parker and Jason Heier (University of Minnesota) for providing the SIL Angiotensin standards.

---

**Supporting Figure 1. Panel A.**
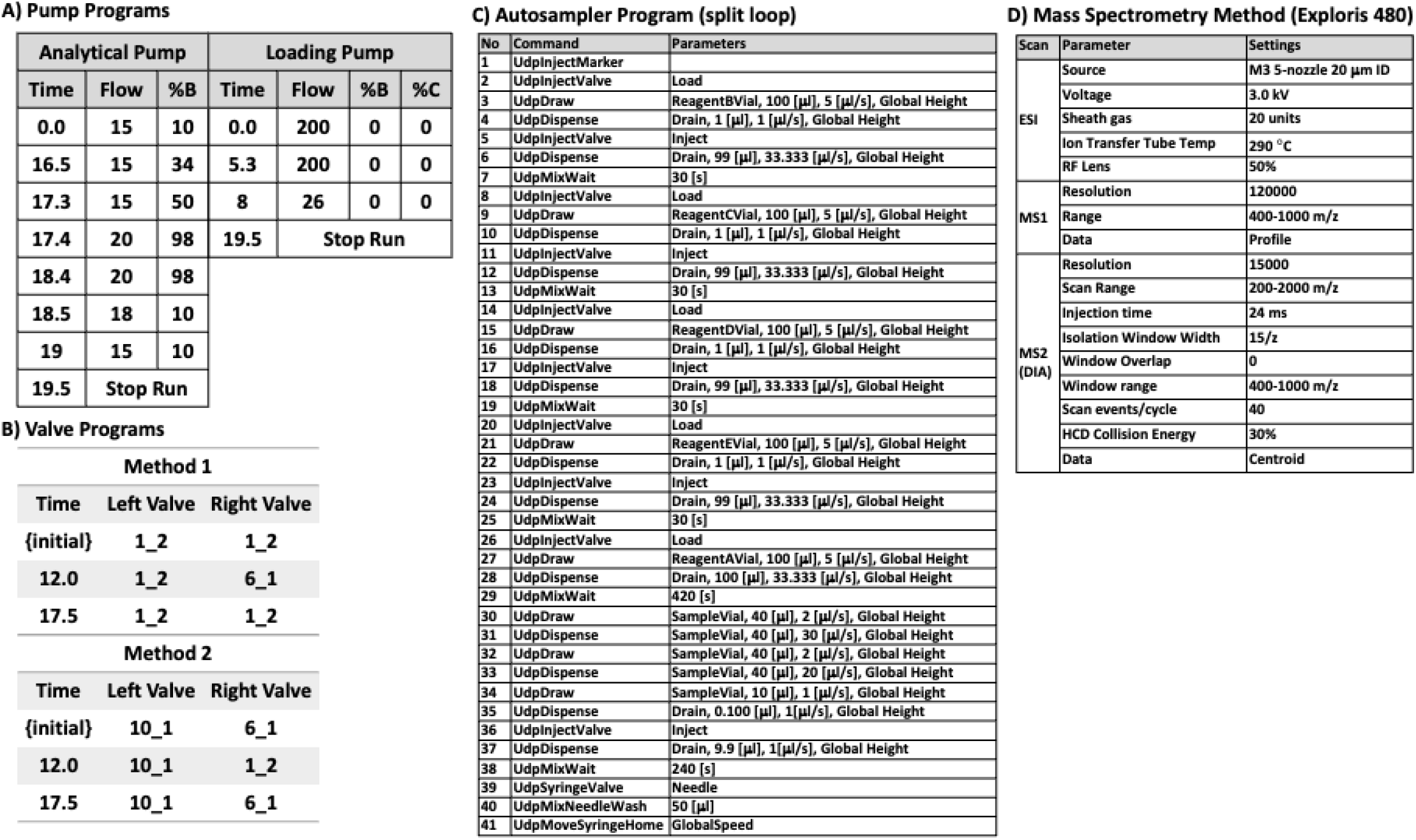
The gradient of the analytical and loading pumps; flow is in μL/min and “Curve” (not shown) was set to 5 for all steps. Analytical pump Buffer B – 0.1% formic acid in 80% acetonitrile, 20% methanol buffer A – 0.1% formic acid in water in both pumps, loading pump Buffers B and C were not used. **Panel B** – The instrument was operated using two alternating methods that only differed in the valve configuration. **Panel C** – Autosampler program, Reagents A and E – water Reagents B and C – methanol, and Reagent D – acetonitrile. **Panel D** – Setting for the mass spectrometry DIA method.

**Supporting Figure 2. Panel A.**
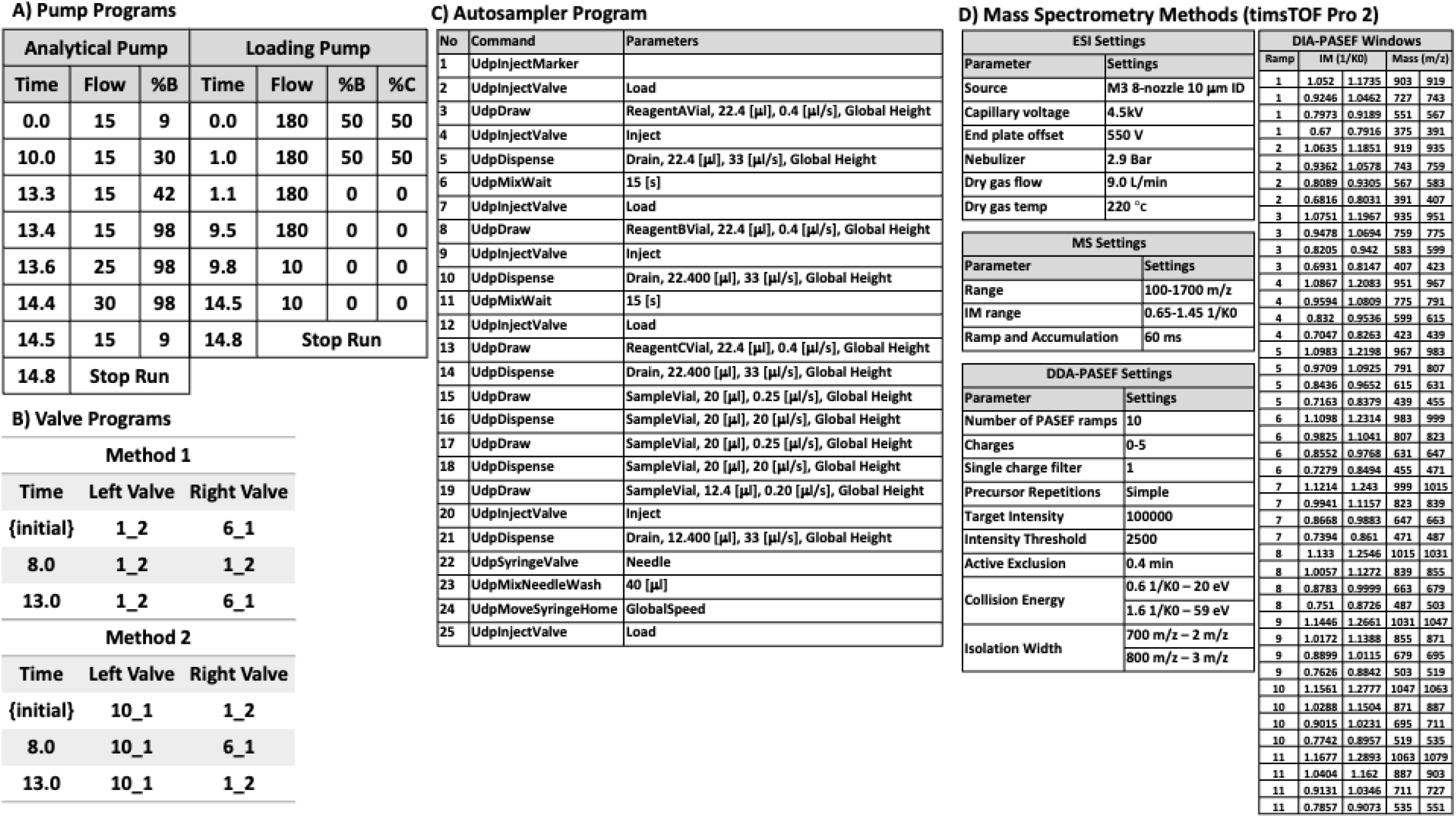
The gradient of the analytical and loading pumps, Analytical pump Buffer B - 0.1% formic acid in 80% acetonitrile, 20% methanol buffer A - 0.1% formic acid in water in both pumps, loading pump Buffer B - 0.1% formic acid in acetonitrile and Buffer C - 100% methanol. **Panel B** – Valve operation for the two alternating methods. **Panel C** – Autosampler program, Reagent A – methanol Reagent B – acetonitrile, and Reagent C – 2% acetonitrile, 0.1% formic acid in water. **Panel D** – Mass spectrometry settings for the DDA and DIA-PASEF methods.

**Supporting Figure 3. Panel A.**
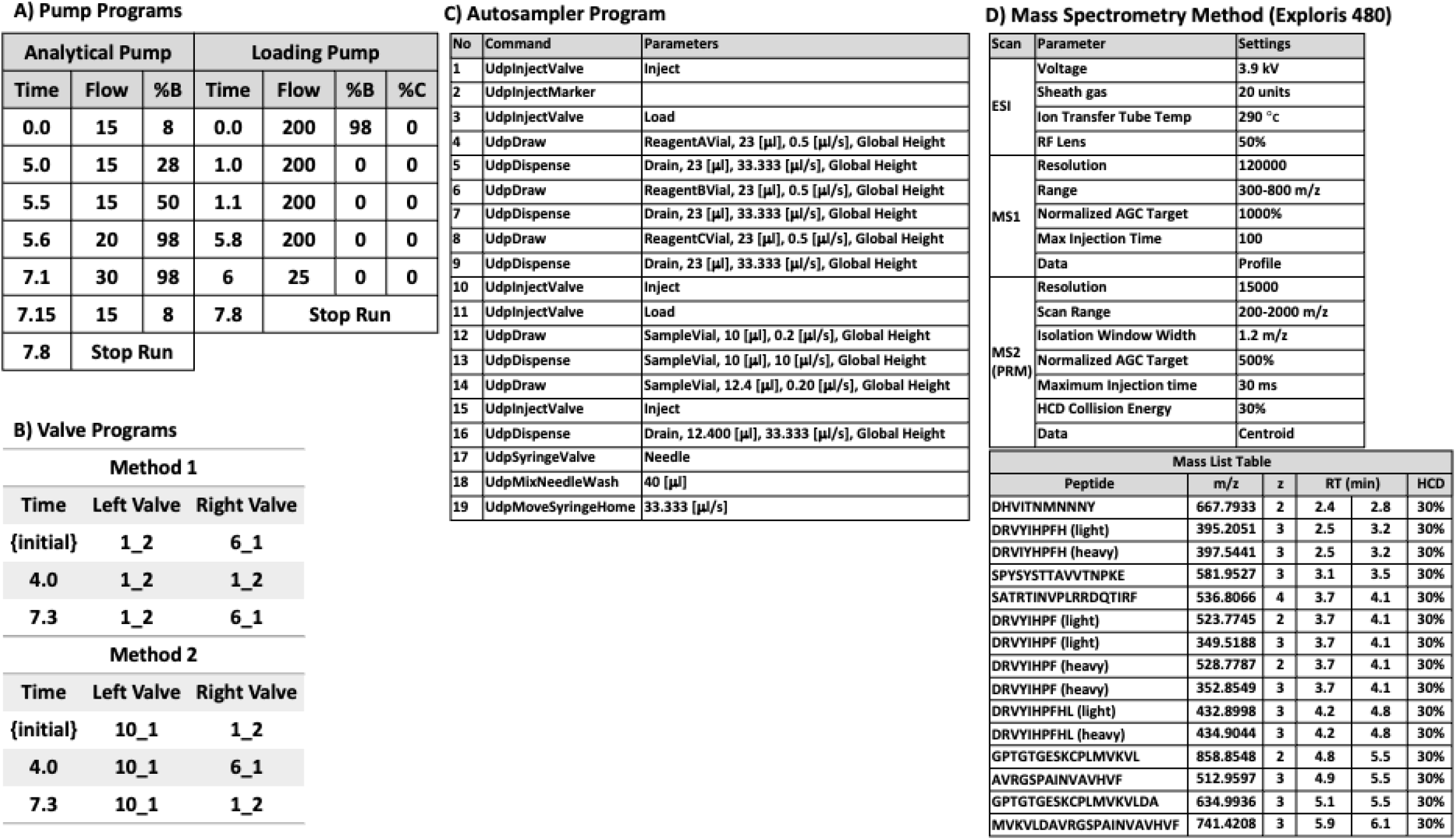
The gradient of the analytical and loading pumps. Analytical pump Buffer B – 0.1% formic acid in 80% acetonitrile, 20% methanol buffer A – 0.1% formic acid in water in both pumps, loading pump Buffer B – 0.1% formic acid in acetonitrile. **Panel B** – Valve operation for the two alternating methods. **Panel C** – Autosampler program, Reagent A – methanol Reagent B – acetonitrile, and Reagent C – 2% acetonitrile, 0.1% formic acid in water. **Panel D** – Mass spectrometry settings for PRM method targeting Angiotensin I and II and peptides from C1Qb and TTR.

## Cited Works

(1) Burkhart, J. M.; Schumbrutzki, C.; Wortelkamp, S.; Sickmann, A.; Zahedi, R. P. Systematic and Quantitative Comparison of Digest Efficiency and Specificity Reveals the Impact of Trypsin Quality on MS-Based Proteomics. J Proteomics 2012, 75 (4), 1454–1462. 10.1016/j.jprot.2011.11.016.

(2) Demichev, V.; Messner, C. B.; Vernardis, S. I.; Lilley, K. S.; Ralser, M. DIA-NN: Neural Networks and Interference Correction Enable Deep Proteome Coverage in High Throughput. Nat Methods 2020, 17 (1), 41–44. 10.1038/s41592-019-0638-x.

(3) Gessulat, S.; Schmidt, T.; Zolg, D. P.; Samaras, P.; Schnatbaum, K.; Zerweck, J.; Knaute, T.; Rechenberger, J.; Delanghe, B.; Huhmer, A.; Reimer, U.; Ehrlich, H. C.; Aiche, S.; Kuster, B.; Wilhelm, M. Prosit: Proteome-Wide Prediction of Peptide Tandem Mass Spectra by Deep Learning. Nat Methods 2019, 16 (6), 509–518. 10.1038/s41592-019-0426-7.

(4) Mansuri, M. S.; Bathla, S.; Lam, T. K. T.; Nairn, A. C.; Williams, K. R. Optimal Conditions for Carrying out Trypsin Digestions on Complex Proteomes: From Bulk Samples to Single Cells. J Proteomics 2024, 297, 105109. 10.1016/J.JPROT.2024.105109.

(5) Vandermarliere, E.; Mueller, M.; Martens, L. Getting Intimate with Trypsin, the Leading Protease in Proteomics. Mass Spectrom Rev 2013, 32 (6), 453–465. 10.1002/mas.21376.

(6) Zougman, A.; Selby, P. J.; Banks, R. E. Suspension Trapping (STrap) Sample Preparation Method for Bottom-up Proteomics Analysis. Proteomics 2014, 14 (9), 1000–1006. 10.1002/pmic.201300553.

(7) Tsai, C. F.; Zhang, P.; Scholten, D.; Martin, K.; Wang, Y. T.; Zhao, R.; Chrisler, W. B.; Patel, D. B.; Dou, M.; Jia, Y.; Reduzzi, C.; Liu, X.; Moore, R. J.; Burnum-Johnson, K. E.; Lin, M. H.; Hsu, C. C.; Jacobs, J. M.; Kagan, J.; Srivastava, S.; Rodland, K. D.; Steven Wiley, H.; Qian, W. J.; Smith, R. D.; Zhu, Y.; Cristofanilli, M.; Liu, T.; Liu, H.; Shi, T. Surfactant-Assisted One-Pot Sample Preparation for Label-Free Single-Cell Proteomics. Commun Biol 2021, 4 (1), 265. 10.1038/s42003-021-01797-9.

(8) Wiśniewski, J. R.; Zougman, A.; Nagaraj, N.; Mann, M. Universal Sample Preparation Method for Proteome Analysis. Nat Methods 2009, 6 (5), 359–362. 10.1038/nmeth.1322.

(9) Giansanti, P.; Tsiatsiani, L.; Low, T. Y.; Heck, A. J. R. Six Alternative Proteases for Mass Spectrometry-Based Proteomics beyond Trypsin. Nat Protoc 2016, 11 (5), 993–1006. 10.1038/nprot.2016.057.

(10) McCabe, M. C.; Gejji, V.; Barnebey, A.; Siuzdak, G.; Hoang, L. T.; Pham, T.; Larson, K. Y.; Saviola, A. J.; Yannone, S. M.; Hansen, K. C. From Volcanoes to the Bench: Advantages of Novel Hyperthermoacidic Archaeal Proteases for Proteomics Workflows. J Proteomics 2023, 289. 10.1016/j.jprot.2023.104992.

(11) Skeggs, L. T.; Kahn, J. R.; Lentz, K.; Shumway, N. P. THE PREPARATION, PURIFICATION, AND AMINO ACID SEQUENCE OF A POLYPEPTIDE RENIN SUBSTRATE.

(12) Harrison-Bernard, L. M. The Renal Renin-Angiotensin System. Adv Physiol Educ 2009, 33 (4), 270–274. 10.1152/advan.00049.2009.

(13) Deng, G. F.; Lin Sun, Y.; Hamet, P.; Inagami, T.; Fu Guo, D. The Angiotensin II Type 1 Receptor and Receptor-Associated Proteins; 2001; Vol. 11. http://www.cell-research.com.

(14) Messerli, F. H.; Bangalore, S.; Bavishi, C.; Rimoldi, S. F. Angiotensin-Converting Enzyme Inhibitors in Hypertension: To Use or Not to Use? Journal of the American College of Cardiology. Elsevier USA April 3, 2018, pp 1474–1482. 10.1016/j.jacc.2018.01.058.

(15) Heil, L. R.; Damoc, E.; Arrey, T. N.; Pashkova, A.; Denisov, E.; Petzoldt, J.; Peterson, A. C.; Hsu, C.; Searle, B. C.; Shulman, N.; Riffle, M.; Connolly, B.; MacLean, B. X.; Remes, P. M.; Senko, M. W.; Stewart, H. I.; Hock, C.; Makarov, A. A.; Hermanson, D.; Zabrouskov, V.; Wu, C. C.; MacCoss, M. J. Evaluating the Performance of the Astral Mass Analyzer for Quantitative Proteomics Using Data-Independent Acquisition. J Proteome Res 2023, 22 (10), 3290–3300. 10.1021/acs.jproteome.3c00357.

(16) Vitko, D.; Chou, W. F.; Nouri Golmaei, S.; Lee, J. Y.; Belthangady, C.; Blume, J.; Chan, J. K.; Flores-Campuzano, G.; Hu, Y.; Liu, M.; Marispini, M. A.; Mora, M. G.; Ramaswamy, S.; Ranjan, P.; Williams, P. .; Zawada, R. J. X.; Ma, P.; Wilcox, B. E. TimsTOF HT Improves Protein Identification and Quantitative Reproducibility for Deep Unbiased Plasma Protein Biomarker Discovery. J Proteome Res 2024, 23 (3), 929–938. 10.1021/acs.jproteome.3c00646.

(17) Guzman, U. H.; Martinez-Val, A.; Ye, Z.; Damoc, E.; Arrey, T. N.; Pashkova, A.; Renuse, S.; Denisov, E.; Petzoldt, J.; Peterson, A. C.; Harking, F.; Østergaard, O.; Rydbirk, R.; Aznar, S.; Stewart, H.; Xuan, Y.; Hermanson, D.; Horning, S.; Hock, C.; Makarov, A.; Zabrouskov, V.; Olsen, J. V. Ultra-Fast Label-Free Quantification and Comprehensive Proteome Coverage with Narrow-Window Data-Independent Acquisition. Nat Biotechnol 2024. 10.1038/s41587-023-02099-7.

(18) Serrano, L. R.; Peters-Clarke, T. M.; Arrey, T. N.; Damoc, N. E.; Robinson, M. L.; Lancaster, N. M.; Shishkova, E.; Moss, C.; Pashkova, A.; Sinitcyn, P.; Brademan, D. R.; Quarmby, S. T.; Peterson, A. C.; Zeller, M.; Hermanson, D.; Stewart, H.; Hock, C.; Makarov, A.; Zabrouskov, V.; Coon, J. J. Journal Pre-Proof The One Hour Human Proteome. Molecular & Cellular Proteomics 2024. 10.1016/j.mcpro.2024.100760.

(19) Kreimer, S.; Binek, A.; Chazarin, B.; Cho, J. H.; Haghani, A.; Hutton, A.; Marban, E.; Mastali, M.; Mesquita, T.; Meyer, J.; Song, Y.; Van Eyk, J.; Parker, S. High-Throughput Single-Cell Proteomic Analysis of Organ-Derived Heterogeneous Cell Populations by Nanoflow Dual-Trap Single-Column Liquid Chromatography. Anal Chem 95 (24), 9145–9150. 10.1021/acs.analchem.3c00213.

(20) Kreimer, S.; Haghani, A.; Binek, A.; Hauspurg, A.; Seyedmohammad, S.; Rivas, A.; Momenzadeh, A.; Meyer, J.; Raedschelders, K.; Van Eyk, J. Parallelization with Dual-Trap Single-Column Configuration Maximizes Throughput of Proteomic Analysis. Anal Chem 94 (36), 12452–12460. 10.1021/acs.analchem.2c02609.

(21) Webber, K. G. I.; Truong, T.; Johnston, S. M.; Zapata, S. E.; Liang, Y.; Davis, J. M.; Buttars, A. D.; Smith, F. B.; Jones, H. E.; Mahoney, A. C.; Carson, R. H.; Nwosu, A. J.; Heninger, J. L.; Liyu, A. V.; Nordin, G. P.; Zhu, Y.; Kelly, R. T. Label-Free Profiling of up to 200 Single-Cell Proteomes per Day Using a Dual-Column Nanoflow Liquid Chromatography Platform. Anal Chem 2022, 94 (15), 6017–6025. 10.1021/acs.analchem.2c00646.

(22) Mc Ardle, A.; Binek, A.; Moradian, A.; Orgel, B. C.; Rivas, A.; Washington, K. E.; Phebus, C.; Manalo, D. M.; Go, J.; Venkatraman, V.; Coutelin Johnson, C. W.; Fu, Q.; Cheng, S.; Raedschelders, K.; Fert-Bober, J.; Pennington, S. R.; Murray, C. I.; Van Eyk, J. E. Standardized Workflow for Precise Mid-And High-Throughput Proteomics of Blood Biofluids. Clin Chem 2022, 68 (3), 450–460. 10.1093/clinchem/hvab202.

(23) Kreimer, S.; Haghani, A.; Binek, A.; Hauspurg, A.; Seyedmohammad, S.; Rivas, A.; Momenzadeh, A.; Meyer, J. G.; Raedschelders, K.; Van Eyk, J. E. Parallelization with Dual-Trap Single-Column Configuration Maximizes Throughput of Proteomic Analysis. Anal Chem 2022, 94 (36), 12452–12460.

(24) Hughes, C. S.; Moggridge, S.; Müller, T.; Sorensen, P. H.; Morin, G. B.; Krijgsveld, J. Single-Pot, Solid-Phase-Enhanced Sample Preparation for Proteomics Experiments. Nat Protoc 2019, 14 (1), 68–85. 10.1038/s41596-018-0082-x.

(25) Hendricks, N. G.; Bhosale, S. D.; Keoseyan, A. J.; Ortiz, J.; Stotland, A.; Seyedmohammad, S.; Nguyen, C. D. L.; Bui, J.; Moradian, A.; Mockus, S. M.; Van Eyk, J. E. An Inflection Point in High-Throughput Proteomics with Orbitrap Astral: Analysis of Biofluids, Cells, and Tissues. 10.1101/2024.04.26.591396.

(26) Kong, A. T.; Leprevost, F. V.; Avtonomov, D. M.; Mellacheruvu, D.; Nesvizhskii, A. I. MSFragger: Ultrafast and Comprehensive Peptide Identification in Mass Spectrometry-Based Proteomics. Nat Methods 2017, 14 (5), 513–520. 10.1038/nmeth.4256.

(27) Yu, F.; Haynes, S. E.; Teo, G. C.; Avtonomov, D. M.; Polasky, D. A.; Nesvizhskii, A. I. Fast Quantitative Analysis of TimsTOF PASEF Data with MSFragger and IonQuant. Molecular and Cellular Proteomics 2020, 19 (9), 1575–1585. 10.1074/mcp.TIR120.002048.

(28) Demichev, V.; Szyrwiel, L.; Yu, F.; Teo, G. C.; Rosenberger, G.; Niewienda, A.; Ludwig, D.; Decker, J.; Kaspar-Schoenefeld, S.; Lilley, K. S.; Mülleder, M.; Nesvizhskii, A. I.; Ralser, M. Dia-PASEF Data Analysis Using FragPipe and DIA-NN for Deep Proteomics of Low Sample Amounts. Nat Commun 2022, 13 (1). 10.1038/s41467-022-31492-0.

(29) Pino, L. K.; Searle, B. C.; Bollinger, J. G.; Nunn, B.; MacLean, B.; MacCoss, M. J. The Skyline Ecosystem: Informatics for Quantitative Mass Spectrometry Proteomics. Mass Spectrom Rev 2020, 39 (3), 229–244. 10.1002/mas.21540.

(30) Jiang, X.; Yeung, D.; Liu, Y.; Spicer, V.; Afshari, H.; Lao, Y.; Lin, F.; Krokhin, O.; Zahedi, R. P. Accelerating Proteomics Using Broad Specificity Proteases. J Proteome Res 2023. 10.1021/acs.jproteome.3c00852.

(31) Zhu, Y.; Wang, Z.; Li, Y.; Peng, H.; Liu, J.; Zhang, J.; Xiao, X. The Role of CREBBP/EP300 and Its Therapeutic Implications in Hematological Malignancies. Cancers. MDPI February 1, 2023. 10.3390/cancers15041219.

(32) Zardab, M.; Stasinos, K.; Grose, R. P.; Kocher, H. M. The Obscure Potential of AHNAK2. Cancers (Basel) 2022, 14 (3), 528. 10.3390/cancers14030528.

(33) Xiong, Y.; Wang, T.; Wang, W.; Zhang, Y.; Zhang, F.; Yuan, J.; Qin, F.; Wang, X. Plasma Proteome Analysis Implicates Novel Proteins as Potential Therapeutic Targets for Chronic Kidney Disease: A Proteome-Wide Association Study. Heliyon 2024, 10 (11). 10.1016/j.heliyon.2024.e31704.

(34) Shen, J.; Zhai, J.; Wu, X.; Xie, G.; Shen, L. Serum Proteome Profiling Reveals SOX3 as a Candidate Prognostic Marker for Gastric Cancer. J Cell Mol Med 2020, 24 (12), 6750–6761. 10.1111/jcmm.15326.

